# Spatiotemporally resolved transcriptomics reveals subcellular RNA kinetic landscape

**DOI:** 10.1101/2022.09.27.509606

**Authors:** Jingyi Ren, Haowen Zhou, Hu Zeng, Connie Kangni Wang, Jiahao Huang, Xiaojie Qiu, Kamal Maher, Zuwan Lin, Yichun He, Xin Tang, Brian Li, Jia Liu, Xiao Wang

## Abstract

Spatiotemporal regulation of the cellular transcriptome is crucial for proper protein expression and cellular function^1^. However, the intricate subcellular dynamics of RNA synthesis, decay, export, and translocation remain obscured due to the limitations of existing transcriptomics methods^2–8^. Here, we report a spatiotemporally resolved RNA mapping method (TEMPOmap) to uncover subcellular RNA profiles across time and space at the single-cell level in heterogeneous cell populations. TEMPOmap integrates pulse-chase metabolic labeling of the transcriptome with highly multiplexed three-dimensional (3D) *in situ* sequencing to simultaneously profile the age and location of individual RNA molecules. Using TEMPOmap, we constructed the subcellular RNA kinetic landscape of 991 genes in human HeLa cells from upstream transcription to downstream subcellular translocation. Clustering analysis of critical RNA kinetic parameters across single cells revealed kinetic gene clusters whose expression patterns were shaped by multi-step kinetic sculpting. Importantly, these kinetic gene clusters are functionally segregated, suggesting that subcellular RNA kinetics are differentially regulated to serve molecular and cellular functions in cell-cycle dependent manner. Together, these single-cell spatiotemporally resolved transcriptomics measurements provide us the gateway to uncover new gene regulation principles and understand how kinetic strategies enable precise RNA expression in time and space.

## Introduction

Cell state and function are shaped by the spatiotemporal regulation of gene expression. This heterogeneous expression is, in part, achieved through precise mRNA metabolism and trafficking over time. The ability to systematically profile transcriptomes across time and space at a single-cell level from intact cellular networks is critical to understanding transcriptional and post-transcriptional gene regulatory mechanisms in cells and tissues.

However, current transcriptomic approaches are unable to simultaneously capture both the spatial and time dependence of RNA profiles. For instance, spatially resolved transcriptomics methods have enabled integrated profiling of gene expression from heterogeneous cell types in the context of tissue morphology^2–8^. Nonetheless, these spatial transcriptomics approaches alone can only provide static snapshots of cells and tissues, while the dynamic flow of gene expression cannot be determined^1^. In contrast, existing metabolic RNA labeling approaches have enabled temporal profiling of the nascent single-cell transcriptome but lack spatial resolution^9–13^. In addition, live-cell imaging can directly track RNA trajectory inside cells, but simultaneously visualizing multiplexed transcripts remains challenging^14^. Thus, there exists a pressing need for highly-multiplexed, spatially and temporally-resolved sequencing methods that tracks nascent mRNAs *in situ* from birth to death at subcellular and single-cell resolutions.

Here, to provide a systematic single-cell analysis of RNA life cycle in time and space, we introduce TEMPOmap (<u>tempo</u>rally resolved *in situ* sequencing and <u>map</u>ping), a method that tracks the spatiotemporal evolution of the nascent transcriptomes over time at subcellular resolution (Extended data Fig. 1a). TEMPOmap integrates metabolic labeling and selective amplification of pulse-labeled nascent transcriptomes with the current state-of-the-art three-dimensional (3D) *in situ* RNA sequencing at 200 nm resolution within a hydrogel-cell scaffold^2^ (Fig. 1a). Using pulse-chase labeling, we were able to simultaneously track key kinetic parameters for hundreds to thousands of genes during their RNA life cycle, including rates of transcription, decay, nuclear export, and cytoplasmic translocation. Using these spatiotemporal parameters, we show that mRNAs of different genes are kinetically sorted at different steps of the RNA life cycle and across different cell-cycle phases, which ultimately serves gene functions.

**Fig. 1.**
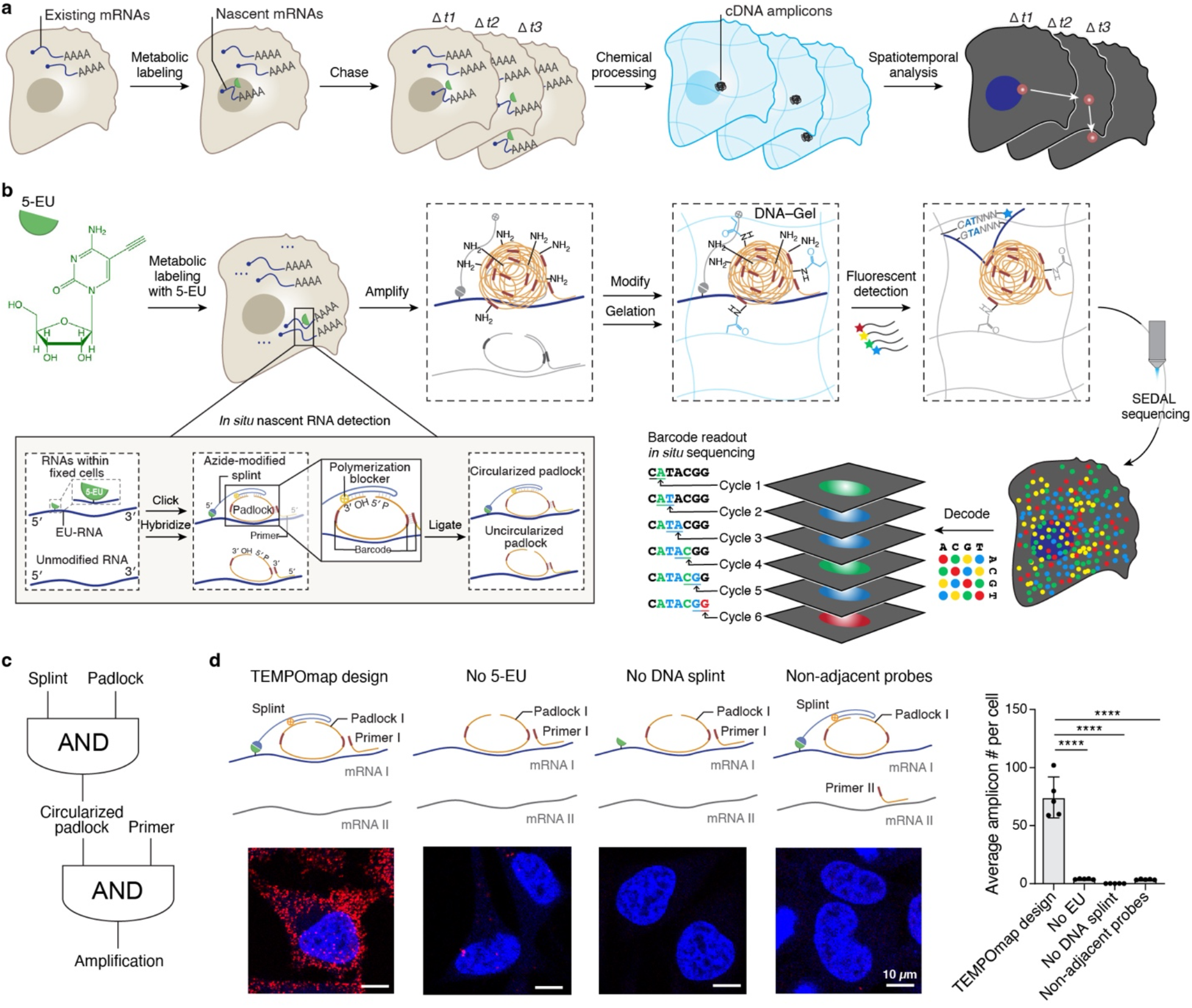
TEMPOmap enables spatiotemporally resolved transcriptomics. **a**, Overview of TEMPOmap pipeline: nascent RNAs of multiple time points are collected and *in situ* sequenced, followed by spatiotemporal RNA analyses. **b**, TEMPOmap experimental workflow. After 5-EU labelled cells are prepared, a set of tri-probes (splint, primer and padlock) are conjugated or hybridized to cellular mRNAs (Extended Data Fig. 1c for more details), resulting in the enzymatic replication of each padlock sequence into cDNA amplicons. The amplicons are anchored *in situ* via a functionalized acrylic group (blue) to a hydrogel mesh to create a DNA-gel hybrid (blue wavy lines). The five-base barcode on each amplicon is read out by six rounds of SEDAL (sequencing with error-reduction by dynamic annealing and ligation). Thus, multiplexed RNA quantification reveals gene expression in nascent subcellular locations. **c**, DNA tri-probe design rationale. The generation of an amplicon requires the presence of splint, circularized padlock, and primer probes in proximity. **d**, Left: schematics and representative fluorescent cell images of negative control experiments of **c**, showing three-part probe requirement for signal amplification. *mRNA_I* represents *ACTB* and *mRNA_II* represents *GAPDH*. All four images show *ACTB* (red) mRNA in HeLa cells (DAPI in blue). Right: quantification of cell images showing the average amplicon reads per cells (*n* = 5 images were measured containing 469, 305, 714 and 520 cells for each condition from left to right, respectively). \*\*\*\**p* <10^−4^, two-tailed *t*-test. Data shown as mean + s.d. Scale bars: 10 *μ*m.

### TEMPOmap strategy for spatiotemporally resolved transcriptomics

TEMPOmap begins with metabolically labeling the cultured cells using 5-ethynyl uridine (5-EU)^13,15^, which adds a bioorthogonal chemical handle on the labeled mRNAs (Fig. 1b). Next, we designed a tri-probe set (splint, padlock, and primer) for each mRNA species to selectively generate complementary DNA (cDNA) amplicons derived from metabolically-labeled RNAs (Fig. 1b-c and Extended data Fig. 1b-c): (1) the splint DNA probe is modified with 5′ azide- and 3′ chain-terminator groups to covalently attach with the 5-EU labeled mRNAs via copper(I)-catalyzed azide-alkyne cycloaddition (CuAAC, Extended data Fig. 1b), thus excluding unlabeled RNAs from subsequent cDNA amplification; (2) the padlock probes recognize mRNA targets with 20-25 nucleotide (nt) cDNA sequence and gene barcodes, which can be circularized when the splint probe is in physical proximity on the same RNA; (3) the primer probes target the neighboring 20-25 nt next to the padlock probes, which serve as the primer to amplify circularized padlocks *in situ* via rolling cycle amplification (RCA), forming cDNA nanoballs (amplicons); in combination, only mRNAs that are bound by all three types of probes will be amplified for selective detection of labeled mRNA population in a label- and sequence-controlled manner via a two-step thresholding strategy (Fig. 1d). Notably, a single gene-targeting padlock probe (bi-probe design) cannot achieve specific gene detection (Extended data Figure 1c) and the dual gene-targeting primer and padlock pair in the tri-probe design is necessary^1^. For proof of concept, we tested representative tri-probes targeted for *ACTB* in HeLa cells, demonstrating specific detection of metabolically labeled transcripts (Fig. 1d, Extended data Fig. 1e). For highly multiplexed transcriptome detection, the *in situ* generated cDNA amplicon libraries are subsequently embedded in a hydrogel matrix for multiple cycles of fluorescent imaging to decode the gene-encoding barcodes via SEDAL (sequencing with error-reduction by dynamic annealing and ligation) (Fig. 1b, Extended data Fig. 1d) to simultaneously detect hundreds to thousands of genes. After the completion of sequencing cycles, the amplicon reads are subsequently registered, decoded, and subjected to 3D segmentation for subcellular and single-cell resolved analysis (Extended data Fig. 2a).

### Spatiotemporal evolution of single-cell nascent transcriptome

To assess TEMPOmap in human cells, we mapped a curated list of 991 genes (981 coding, 10 non-coding RNA) with diversified spatial and temporal RNA expression profiles^13,16^ in HeLa cell cultures. Then, we designed a pulse-chase experiment^13,17^ with one hour (hr) of pulse labeling and various chase times (0, 1, 2, 4, and 6 hrs) as well as one steady-state reference with 20-hour pulse labeling (Fig. 2a), followed by the TEMPOmap experiment workflow (Fig. 1b). The barcodes in all the samples were sequenced over six rounds of *in situ* sequencing, followed by a final round of subcellular compartment staining (nuclei and cytoplasm) to segment cell bodies and assign the subcellular locations of amplicons in 19,856 cells in 3D (Extended data Fig. 2b-d). From 0 to 6 hrs chase time post-labeling, we observed a decline of total RNA reads per cell, a gradual shift of the RNA distribution from the nucleus to the cytoplasm, and further allocation from the middle cytoplasmic region to the periphery (Fig. 2c, d), in agreement with the expected trajectory of RNAs. Interestingly, a significant fraction of reads (~40%) was retained in the nucleus even after 6 hrs chase. A closer inspection of the retained RNA molecules revealed that RNAs with the highest nuclear-to-cytoplasm read ratio included long non-coding RNAs (*NEAT1, MALAT1*), supported by deep sequencing of RNA from cellular fractions (Extended data Fig. 3a)^18,19^. Notably, a group of mRNAs (*e.g. KIF13A, LENG8, CCNL2, COL7A*) showed high ratio of nuclear retention (nuc/cyto > 2, Extended data Fig. 3a). Our observation validates the previous discovery of nuclear retention of mRNA, which may serve as a regulatory role to buffer the cytoplasmic gene expression noise^20,21^.

**Fig. 2.**
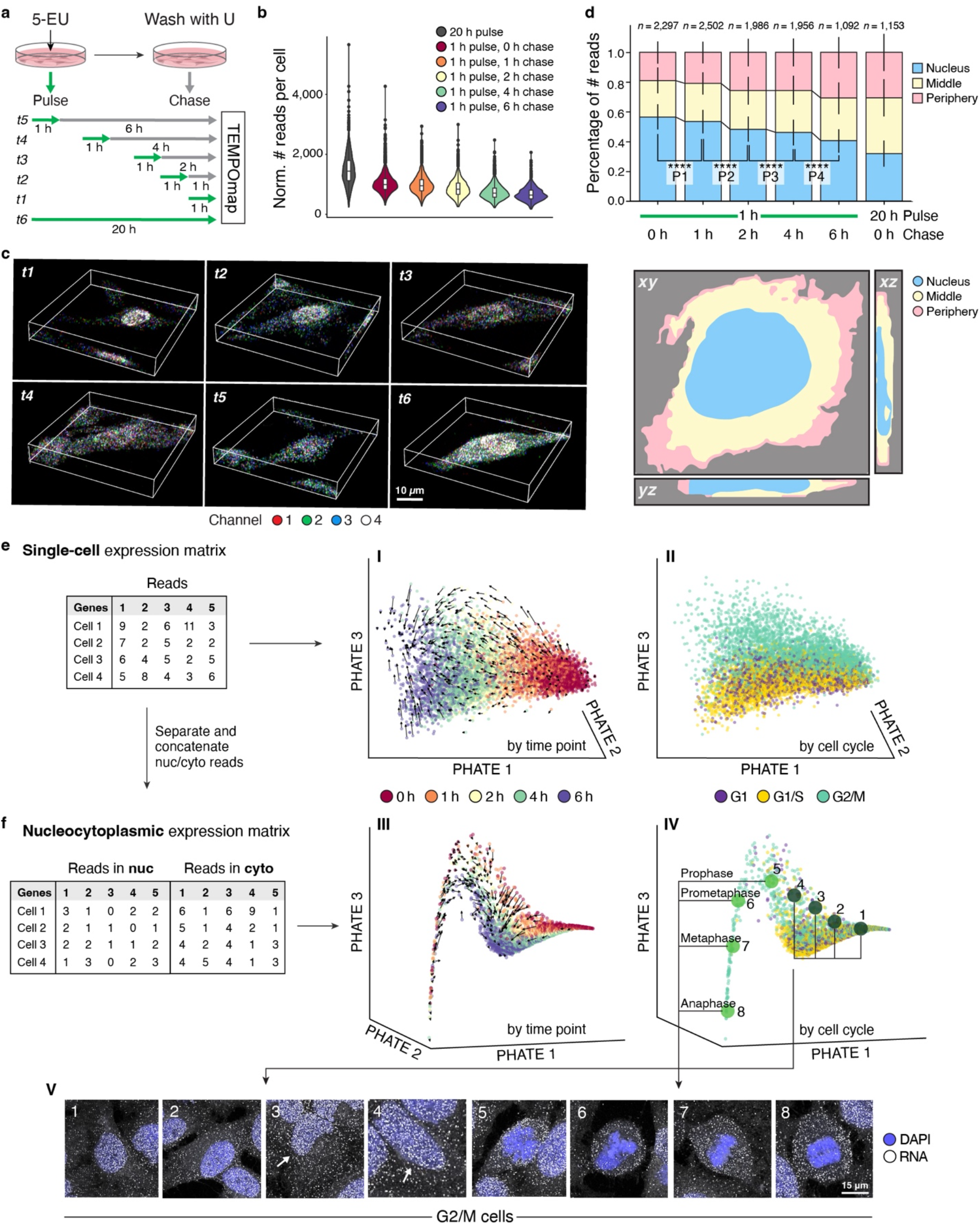
Spatiotemporal tracing of single-cell transcriptome. **a**, Pulse-chase experiment design on HeLa cells. For the first five time points, we used 1 hr metabolic labeling (pulse) followed by 0, 1, 2, 4 and 6 hrs chase. At the last time point, we metabolically labeled the cells for 20 hrs. All of the cells were then processed by TEMPOmap workflow measuring 998 genes. **b**, RNA reads (cDNA amplicons) per cell for each pulse-chase time point. **c**, 3D fluorescent images of in-process TEMPOmap with zoomed views of representative single cells of cycle 1 at each time point. Z-stack range: 10 μm. **d**, Top, boxplot summarizing the fraction of reads in each subcellular region of all cells at each time point. Vertical lines indicate s.d. The statistics compares the fractions of nuclear reads (blue) across the first five time points. \*\*\*\**p* < 0.001, Kruskal–Wallis test with post hoc Tukey’s HSD. Number of cells (n) in each time point is shown. Bottom, subcellular region assignment (nuclear, middle and periphery) of one representative cell. **e-f**, TEMPOmap single-cell (**e**) or nucleocytoplasmic (**f**) RNA measurements rendered as a visualization by PHATE and colored by pulse-chase time points (I, III) or cell-cycle marker gene expression (II, IV). Black arrows inferred by RNA degradation vectors indicate the directions of chase time progression. Bottom row, representative raw images of G2/M phase cells separated on PHATE coordinates. All images show mRNAs (in white) in HeLa cells (DAPI in blue). Scale bars: 15 *μ*m.

Next, we asked whether the TEMPOmap dataset could resolve the heterogeneity of single cells. To this end, we pooled all the cells under the 1 hr pulse conditions (18,176 cells) for single-cell resolved dynamic trajectory analysis using PHATE (Fig. 2e, I)^22,23^. Our results showed a clear trajectory along the progression of chase time, which suggests that the temporally resolved single-cell transcriptional states could be readily distinguished and aligned in the latent space. Overlaying the same coordinates with RNA degradation kinetics vectors (represented as the quivers) further recapitulated the single-cell trajectory along RNA life cycle progression^23–25^. We further asked how the RNA life cycle defined by the pulse-chase timeline aligns with cell-cycle progression. To this end, we classified the cells into three cell cycle phases (G1, G1/S, and G2/M) based on their nascent expression of marker genes (Extended data Fig. 3b,c) using cell-cycle scoring^26^. Interestingly, the direction of cell-cycle progression is orthogonal to that of the pulse-chase time point progression (Fig. 2e, II). This observation suggests that TEMPOmap provided independent temporal information regarding the RNA life cycle in addition to the cell cycle.

Besides single-cell analysis, we considered that TEMPOmap dataset could reveal the subcellular dynamics. To this end, we generated a nucleocytoplasmic gene-by-cell matrix by concatenating single-cell nuclear expression with cytoplasmic expression for trajectory analysis (Fig. 2f). Apart from recovering the unidirectional trajectory of single cells along with the labeling time points (Fig. 2f, III), we found a small fraction (*n* = 137 cells, 2.1%) of G2/M cells formed a narrow trajectory and projected into a distinct space, suggesting that the nucleocytoplasmic RNA distribution in this group of G2/M cells drastically differs from the rest of the G2/M cells (Extended data Fig. 3d). We suspected that these spatially distinct cells were the cells undergoing mitosis with their unique RNA nucleocytoplasmic distribution^27^. Indeed, the cells on this trajectory had been in different phases of mitosis, during which RNAs were mostly evicted from the chromatin regions compared to that in G2 cells (Fig. 2f, V). Furthermore, the uniform direction of this distinct trajectory aligns well with the time progression of mitosis (Fig. 2f, V, 5-8), indicating that the temporal mitotic transitions could be inferred by subcellular RNA localization patterns. As a result, by jointly making use of the time-gated nucleocytoplasmic distribution, we not only separated G2 and M cells but also traced the trajectory of mitosis on the gene expression space, during which M cells undergo drastic RNA eviction from chromosomes^28^.

### Subcellular RNA kinetic landscape across RNA lifespan

To further quantify the kinetics during different stages of transcription and post-transcriptional processing, we estimated four key kinetic constants for all detected transcripts across RNA lifespan – synthesis (α), degradation (β), nuclear export (λ) (Fig. 3a), and cytoplasmic translocation (γ) (Fig. 3b). We noticed a correlated relation between physical cell volumes and single-cell RNA reads (Extended data Fig. 4a-b). To remove the potential bias caused by cell volume, we estimated α and β values based on the averaged concentrations of each RNA species (reads/voxel) across single cells (Extended data Fig. 4c). Built on the previous studies^18,22,30^, our model assumed zero-order kinetics for α and first-order kinetics for β^17,29^. In addition, a threshold of fitting β for each gene (936 genes out of 991 genes with the coefficient of determination *R^2^* >= 0.5) was applied for quality control purposes (Extended data Fig. 4c-d). In parallel, we estimated the nuclear export rate (λ) based on the change in the ratios of nuclear-to-total reads over time. We noted that the estimation of λ might also be complicated by nuclear and cytoplasmic degradation, and therefore was more fitting for describing the change in the homeostasis of nucleocytoplasmic RNA distribution. Lastly, to systematically evaluate the relative positions of each RNA species in physical cytoplasm space in 3D over time, we derived a distance-ratio (DR) based method (Extended data Fig. 2d, Extended data Fig. 4c), where the cytoplasmic translocation rate (γ) was calculated by tracking the change of DR over time (Fig. 3b).

**Fig. 3.**
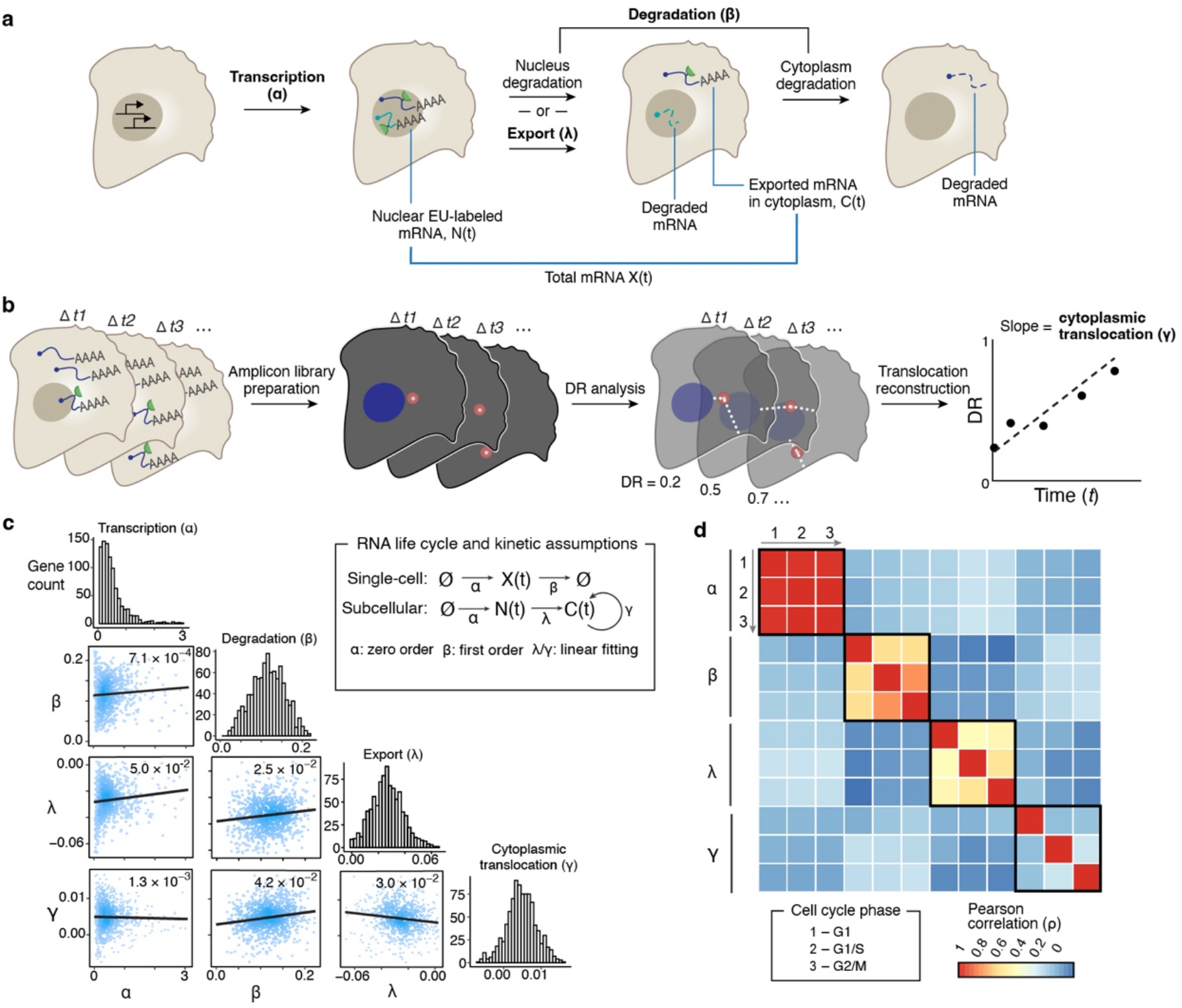
TEMPOmap reveals subcellular RNA kinetic landscape across RNA lifespan and cell cycle. **a**, The dynamic model for estimating RNA kinetic parameters. For each gene, RNA synthesis (α) and degradation constant (β) were estimated using single-cell RNA concentration. The export constant (λ) was estimated using the subcellular RNA concentrations. **b**, The dynamic model for estimating cytoplasmic translocation (γ) using distance ratio (DR)-based analysis (see Methods). **c**, Upper Right: the mathematical model of RNA life cycle and kinetic assumptions used for the parameter estimation. Bottom Left: The histogram of the four parameters for all genes that passed qualify control and the scatter plots depicting the pairwise correlation of parameters with ρ value (Pearson correlation) and linear fitting curve. Color intensity of the dots indicates local density. **d**, Heatmap depicting pairwise correlation matrix of the four parameters estimated using single-cells from three cell-cycle phases (G1, G1/S, G2/M). Color indicates the value of Pearson correlation coefficients. Boxed regions indicate the correlations of each parameter among three cell-cycle phases.

Notably, while nuclear export of RNA had been considered to be a constant in the previous RNA velocity-based model^29^, our result suggested that λ varies substantially among different RNA species, which indicates gene-specific regulatory mechanisms to control the homeostasis of nucleocytoplasmic transcript distribution (Extended data Fig. 4e). In addition, for the first time to the best of our knowledge, we could systematically study the cytoplasmic translocation of RNAs of a large number of genes simultaneously at 1 hr resolution. Most genes had γ > 0 (Extended data Fig. 4f, g), which suggested a translocating direction from the nuclear membrane to the cytoplasmic membrane. However, we found a small subset of genes with γ < 0 (*R^2^* > 0.5) that were significantly enriched in secreted and organellar proteins (Extended data Fig. 4h), indicating possible relocation events from the cytosol to the endoplasmic reticulum or faster degradation rates for non-ER anchored RNAs than ER-anchored ones. Further studies need to be conducted to investigate the kinetic mechanism that directs the cytoplasmic translocation of different RNA molecules (Extended data Fig. 4i).

Next, we asked whether any of the four RNA kinetic parameters were intrinsically coupled. Here, we performed pairwise correlations of the four parameters across 936 genes. We found that the overall correlation between each pair of parameters was weak (Fig. 3c, *ρ* < 0.1), suggesting that the kinetic parameters of RNA transcription, post-transcriptional processing^13^ and allocation are relatively independent^16^. We then explored the correlations of these kinetic parameters across the cell cycle. To this end, we performed a further pairwise correlation analysis of the four parameters across different genes at three cell-cycle phases (800 genes passed quality control; Fig. 3d). Interestingly, for each parameter, depending on its temporal sequence in the RNA life cycle, a trend of decreasing correlations in cell cycle phases emerged: at the early stage of RNA production, the synthesis rates α were highly correlated (*ρ* = 0.9-1.0, Extended data Fig. 5a); during post-transcriptional processing in the nucleus, λ in the three phases have moderate correlations (*ρ* = 0.4-0.5, Extended data Fig. 5c); near the end of the RNA life cycle, cytoplasmic translocation γ have much weaker correlations (*ρ* = 0-0.2, Extended data Fig. 5d). This observation suggested that RNA metabolism and trafficking of different genes become less synchronized and increasingly heterogeneous from the upstream to the downstream stages of RNA life cycle, potentially due to gene-specific and cell-cycle-dependent regulation.

Given the cell-cycle resolved RNA kinetic landscape, we further investigated how RNAs could be dynamically “sculpted” to fine-tune the temporal RNA expression profiles. First, we identified potentially co-regulated RNAs through a pairwise single-cell covariation analysis of 936 genes from the aforementioned pulse-chase HeLa cell samples (1 hr pulse, 0-6 hrs chase, Extended data Fig. 6a, left). Using the matrix of pairwise correlation single-cell expression variation combining all time points, we identified four groups of genes with significant intra-group correlation, indicating potential gene co-regulation patterns (Extended data Fig. 6a, right, Group 1-4). Notably, while these genes are enriched with cell-cycle-related functions (Extended data Fig. 6b), the four groups differ significantly in multiple stages of RNA kinetics (Extended data Fig. 6c). Next, we repeated the single-cell covariation analysis to each individual time point using the same gene order, and found that the shift in the co-variation pattern of each group varies from 0 to 6 hrs (Extended data Fig. 6d): Group 1 shows decreasing co-variation pattern from 0 to 2 hrs post-synthesis; Group 2 shows consistently high expression co-variation across time; in contrast, the co-variation patterns of Group 3 and 4 gradually emerged from 2 hrs to 6 hrs post-synthesis. This observation suggests that, at the RNA level, cell cycle progression is jointly shaped by an orchestration of genes with distinct transcriptional and post-transcriptional kinetic features.

### Differential RNA kinetic strategies by gene function

After recognizing the aforementioned four gene groups whose RNA temporal profiles coupled with cell-cycle phasing, we asked if such correspondence between RNA kinetics and gene functions globally exists for other genes. To identify gene modules based on their shared kinetic patterns in the context of RNA life cycle and cell cycle, we first clustered 800 genes using the 12 parameters (four kinetic constants across three cell cycle stages. The clustering analysis revealed five kinetic gene clusters of distinct kinetic landscapes (Fig. 4a) that also had distinct subcellular distributions over time (Extended data Fig. 7a, b). Importantly, gene ontology analyses showed that the five clusters associate with distinct biological and molecular functions (Fig. 4b). For example, genes with unstable and slowly exported RNAs were strongly enriched in metal-binding and transcription factor binding activities (Cluster 1, *n* = 231 genes); genes with high RNA stability and moderate export rate (Cluster 3, *n* = 153 genes) were enriched in hydrolase and ATP-binding activities. On the other hand, genes with fast synthesis and greater RNA stability (Cluster 5, *n* = 86 genes) were enriched in constitutive cellular processes like mRNA splicing, translation, and mitochondrial functions. We reasoned that these housekeeping genes tend to produce abundant and stable RNAs for a longer persistence of genetic information due to energy cost of protein production^30^.

**Fig. 4.**
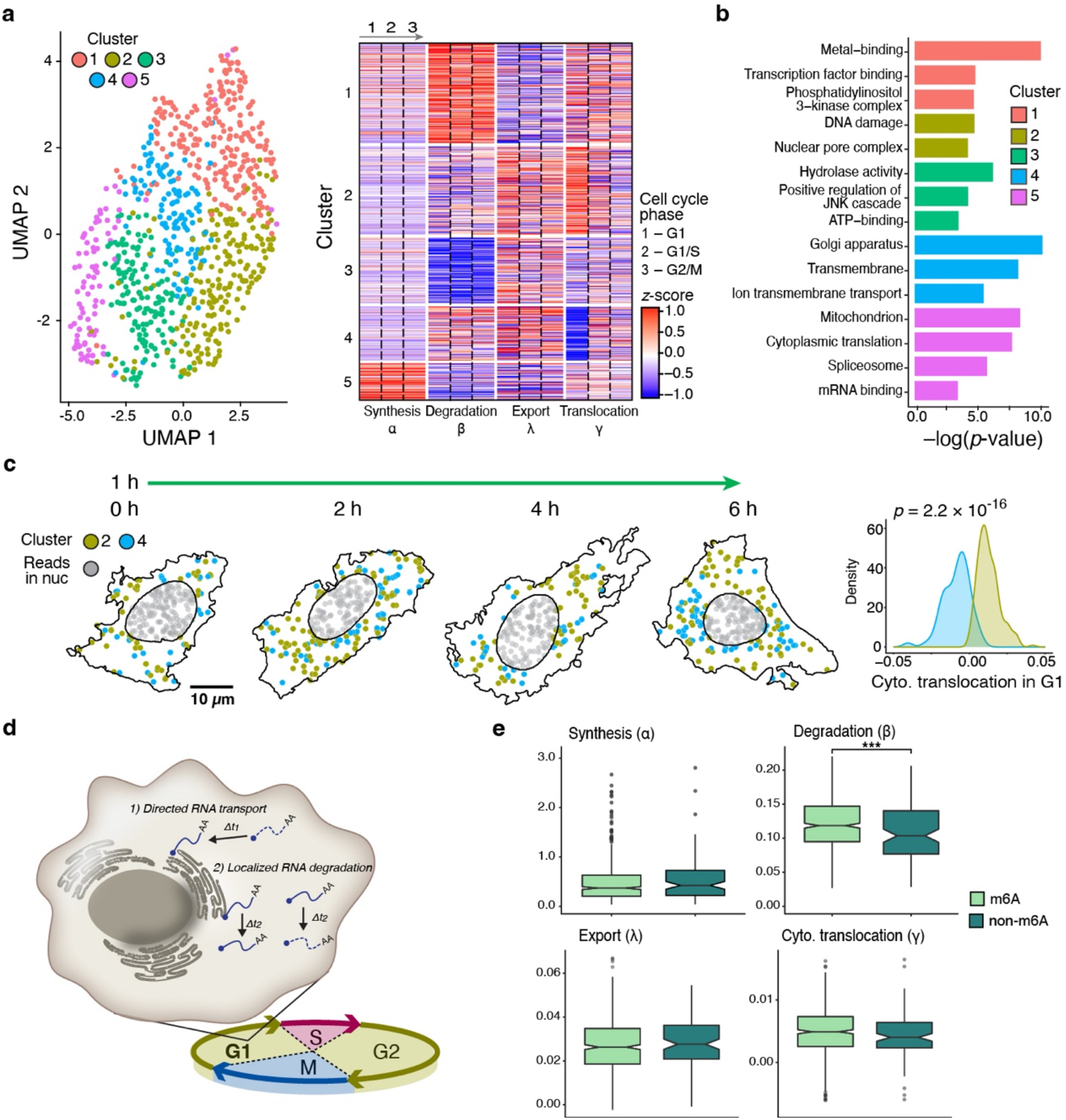
Differential RNA dynamics by gene function and post-transcriptional characteristics. **a**, UMAP representation (left) and heatmap (right) showing the gene clustering using all 18 estimated parameters across cell cycle. Color in the heatmap represents the parameter-wise z-score normalized value. **b**, Pathway enrichment analysis of genes in each cluster in **a** using DAVID. **c**, Left: visualization of cytoplasmic RNAs of Cluster 2 and 4 in representative cells across pulse-chase time points. Scale bar: 10 *μ*m. Right: density plot showing the distributions of γ values of genes in cluster 2 and 4. **d**, diagram illustrating two possible mechanisms of reverse mRNA translocation (γ) at the G1 phase of cluster 4 genes: directed RNA transport and localized RNA degradation. e, Boxplots comparing the four parameters estimated for m^6^A and non-m^6^A genes. Data shown as means (notches), 25-75% quartiles (boxes) and ranges (vertical lines). *** *p*<0.01, Wilcoxin test in **c** and **e**.

Notably, while Cluster 2 (*n* = 205 genes) and 4 (*n* = 125 genes) have slower synthesis, moderate degradation, and faster export, they significantly differ in cytoplasmic translocation rates (γ), which are cell-cycle-dependent. In G1 phase, RNAs of Cluster 2 exhibited significantly higher γ compared to the other phases, whereas Cluster 4 showed the opposite trend (Fig. 4a, right). We found the genes in Cluster 2 have a functional enrichment of DNA damage and repair, and those in Cluster 4 are functionally related to organellar and membrane-bound proteins (Fig. 4b). Closer examination of the genes in Cluster 4 showed that, most of the genes (109/125) have negative γ values in G1, indicating an overall reverse direction of translocation in G1 phase. Previous research showed that many mRNAs encoding membrane-bound proteins were anchored to the surface of endoplasmic reticulum (ER) for localized protein synthesis^31^. We reasoned that the RNAs encoding these membrane proteins might also be regulated at the dynamic level, executed by both spatial and temporal localization control in a cell-cycle-dependent manner. Since the duplication of organelles and cell expansion are the major activities at G1 phase, our discovery also suggests that, ER-localized protein synthesis might be more active in G1, either by RNA transport toward ER or local degradation of non-ER-anchored RNAs in the cell periphery (Fig. 4d). Hence, we proposed a more comprehensive picture of the regulation dynamics of membrane protein at the RNA processing level from both spatial and temporal perspectives. While the mechanism that underlies our observed translocation values is still open to further investigations, we revealed the importance of regulating the spatiotemporal localization of transcripts that carry different genetic information.

Finally, we examined RNA kinetic landscape in the context of *N^6^*-methyladenosine modifications (m^6^A), a critical posttranscriptional chemical modification of RNA that plays vital physiological roles^32,33^. RNA methylation m^6^A is known to mediate a wide range of post-transcriptional gene regulation, however, the full landscape of the spatiotemporal dynamics on m^6^A-RNA has not been systematically addressed. To this end, we separated the genes encoding RNAs with and without m^6^A modifications by previous m^6^A profiling studies^34,35^ (m^6^A- or non-m^6^A-RNAs, Extended data Fig. 8a). Consistent with the previous report, m^6^A-modified RNAs were significantly less stable than non-m^6^A-RNAs (higher β, Fig. 4e). In addition, we observed the same trend when comparing the degradation constants in different cell cycle phases, suggesting that regulating the decay of m^6^A-methylated RNA is persistent across cell cycle (Extended data Fig. 8b). Together, we demonstrate the potential of using TEMPOmap dataset to study spatiotemporal transcriptomics in combination with post-transcriptional modification, a path to incorporate multi-modality transcriptomic analysis at single-cell and subcellular resolution.

## Discussion

TEMPOmap serves as a novel *in situ* transcriptomic platform that simultaneously profiles time- and space-resolved transcriptomics in single cells, a multimodal single-cell transcriptomics technology at the subcellular resolution that has not been achieved before. We demonstrated the capacity of TEMPOmap to systematically detect the subcellular allocation and cytoplasmic translocation of transcripts over time. More importantly, our study provided a full landscape of RNA subcellular kinetics at the single-cell level and revealed how RNA kinetics contribute to cellular functions such as cell-cycle progression. We observed a strong correlation of RNA kinetic patterns with the molecular functions of genes-- such function-oriented regulation of RNA life cycle might have evolved under survival and energy constraints to control spatiotemporal gene expression in a precise and economic way^30^. In future work, TEMPOmap can be combined with high-throughput single-cell functional genomics (*e.g*. CRISPR screens^36^) to determine key molecular factors that impact the kinetic landscape of RNA life cycle. Furthermore, such spatiotemporally coordinated transcriptomic patterning may shed light on understanding the molecular mechanisms of various biological phenomena, including in development and pattern formation, learning and memory, biological clocks, as well as disease progression. With optimization of metabolic labeling conditions^15,37,38^ and integration of various molecular probing schemes, such methodology can be adapted for *ex vivo* or *in vivo* tissue samples to systematically profile dynamic events in tissue biology.

## Acknowledgment

We thank Albert Liu, Jiakun Tian and Jianting Guo for their helpful comments on the manuscript. We also thank Leslie Gaffney for her help in graphic illustration. X.W. acknowledges the support from Searle Scholars Program, Thomas D. and Virginia W. Cabot Professorship, Edward Scolnick Professorship, Ono Pharma Breakthrough Science Initiative Award, Merkin Institute Fellowship, and NIH DP2 New Innovator Award.

## Author contributions

X.W. and J.R. conceived the idea. X.W., J.R., and H.Zeng developed the methodology for the study. J.R. and H.Zeng carried out experimental work. H. Zhou, J.H, K.W, X.Q, Y.H, X.T performed computational and data analyses. Z.L. helped with TEMPOmap data collection. J.R. and X.W. prepared the manuscript. H.Zeng. provided critical discussions during the whole development. X.W., H.Zeng, and H.Zhou critically revised the manuscript. X.W. supervised the study.

---

**Extended Data Fig. 1.**
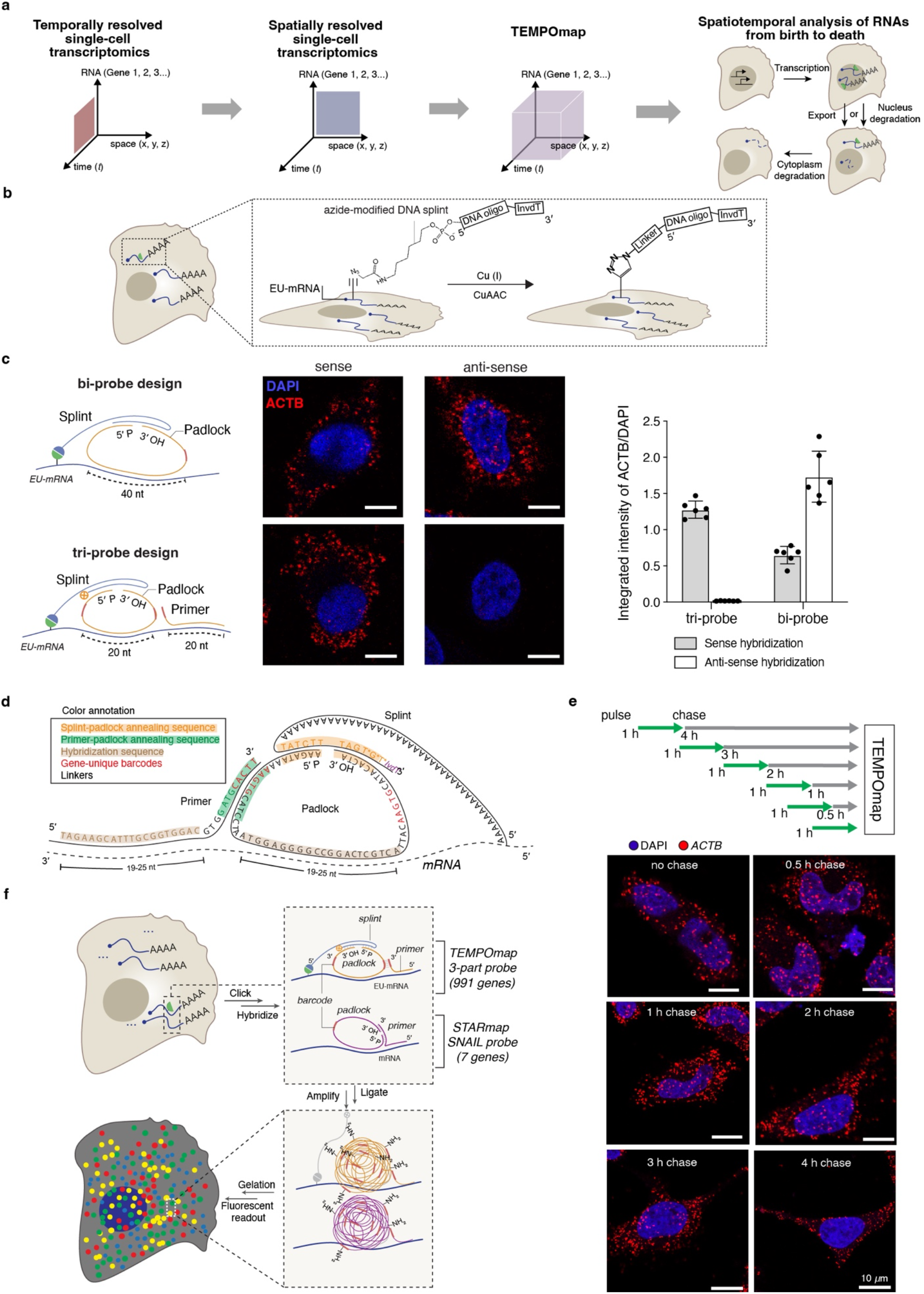
TEMPOmap experimental design and optimization. **a**, Method conceptualization. TEMPOmap combines RNA metabolic labeling and state-of-the-art spatial transcriptomics to achieve single-cell spatiotemporal transcriptomics for RNA dynamic analysis. **b**, CuAAC-mediated click chemistry to conjugate azide-modified splint and EU-labeled nascent transcript. **c**, Comparison of TEMPOmap bi-probe and tri-probe design targeting *ACTB* mRNA. Left, probe design schematics. Middle, representative fluorescent images of cells treated with sense-targeting and antisense-targeting padlocks and primers. Right, quantification of fluorescence in cell images (6 images containing 400-600 cells were measured under each condition). Data shown as mean + s.d. **d**, DNA sequences of TEMPOmap tri-probe system. **e**, Proof-of-concept pulse-chase experiment (top) followed by raw cell images (bottom) showing the translocation of *ACTB* mRNAs when chased after 1 hr EU treatment with different times. Cell nuclei (blue), amplicons (red). **f**, Simultaneous mapping and sequencing of nascent RNAs by TEMPOmap and total RNAs by STARmap in the experimental workflow. TEMPOmap-targeted amplicon reads were normalized against the reads of STARmap-targeted RNAs. Scale bar: 10 *μ*m.

**Extended data Fig. 2.**
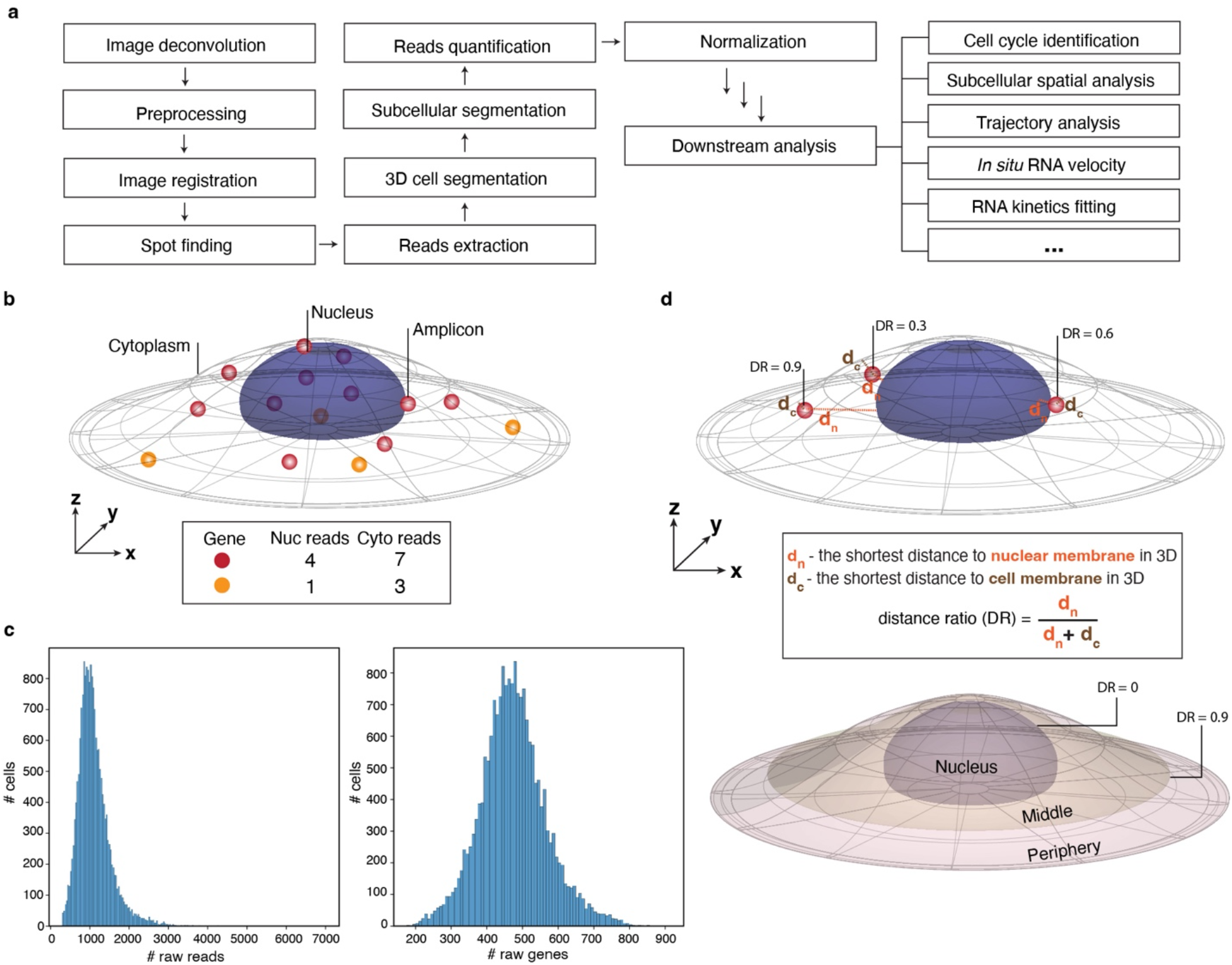
TEMPOmap data processing and analysis. **a**, TEMPOmap data analysis pipeline. **b**, Schematics of reads assignment in subcellular compartments. **c**, Histograms showing detected reads (cDNA amplicons) per cell (left), and genes per cell (right). **d**, Schematics of distance ratio (DR)-based subcellular segmentation in the cytoplasm. Two values for each amplicon were computed in 3D: d_n_, the shortest distance to nuclear membrane; d_c_, the shortest distance to cell membrane. “Middle” is the region defined between DR = 0 and 0.9. “Periphery” is defined as DR > 0.9.

**Extended data Fig. 3.**
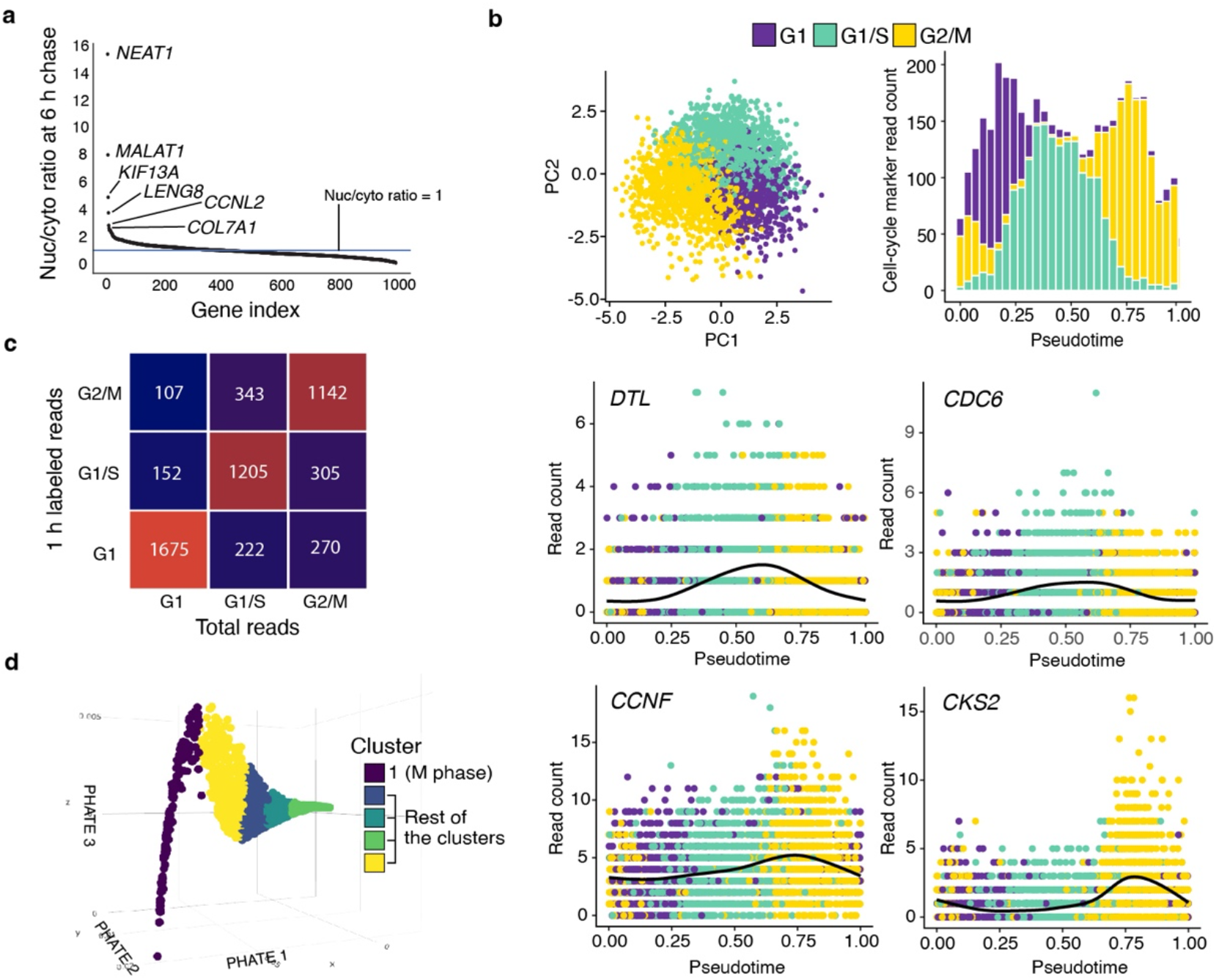
RNA subcellular analysis and cell-cycle phase identification. **a**, nuclear-to-cytoplasmic ratio of amplicon reads of 991 genes at 6 hrs chase time point. Genes were ranked from top to bottom according to the ratios. **b**, Cell-cycle identification (G1, G1/S, G2/M) by cell-cycle gene marker measured via TEMPOmap labeled RNA expression. Cell-cycle scores were calculated via ‘score_genes_cell_cycle’ in scanpy. The cells were visualized via PCA and colored by cell-cycle phases (top). Variations in the raw counts of all cell-cycle gene markers (bottom left) and four representative markers (bottom) were projected by the pseudotime analysis. **c**, Comparison of cell-cycle identification by 1 hr pulse-labeled reads and total reads using scEU-seq dataset^14^ shows that the nascent transcriptome can accurately define cell-cycle states. The number in each box indicates the number of cells. **d**, Cell clustering result based on PHATE embedding of the nucleocytoplasmic matrix. Cluster 1 incorporates the cells (*n* = 137 cells) in the M phase by visual inspection of raw images.

**Extended data Fig. 4.**
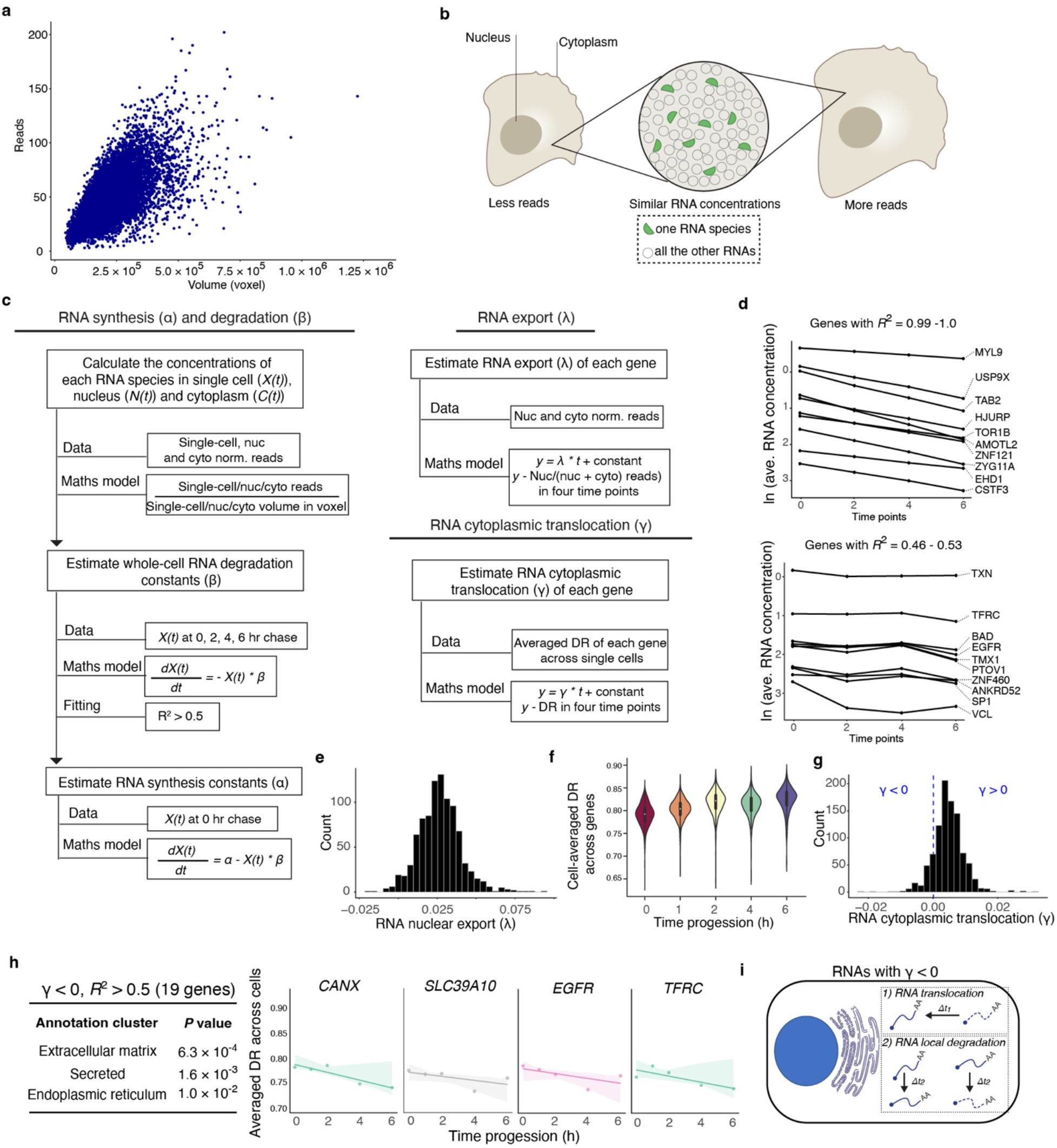
Quantification of RNA subcellular kinetic parameters. **a**, Correlated relation between cell volume (in voxels) and single-cell reads, indicating the influence of transcript number by cell volume. **b**, Schematics showing the biased single-cell RNA read counts with varying physical cell volumes. **c**, Mathematical models for estimating RNA kinetic parameters (α,β and λ) and the detailed workflow of calculation and fitting procedure. Note: *X(t)* = single-cell RNA concentration; *N(t)* = nuclear RNA concentration; *C(t)* = cytoplasmic RNA concentration. **d**, Changes in the natural log of *X(t)* across time points of genes with R^2^ = 0.99-1.0 (top) and R^2^ = 0.46-0.53 (down). Representative genes are shown. The estimated β values for all genes were filtered with a threshold of R^2^ > 0.5 as a quality control. **e**, Histogram of estimated λ (nuclear export) values for all genes. **f**, The distribution of single cell-averaged DR values for all 991 genes across 0-6 hrs chase time points. **g**, Histogram of estimated γ (cytoplasmic translocation) values for all genes. Blue dashed line separates the genes with γ > 0 and γ < 0, which indicates the opposite direction of observed translocation. **h**, Left, 19 genes with γ < 0 (R^2^ > 0.5) were strongly enriched in secreted and organellar proteins. Middle, time-lapsed DR values of representative genes. **i**, Schematics showing the observed inward direction of RNA translocation of genes with γ < 0. Two potential mechanisms are shown: 1). RNA translocation (active or passive); 2). Local RNA degradation.

**Extended data Fig. 5.**
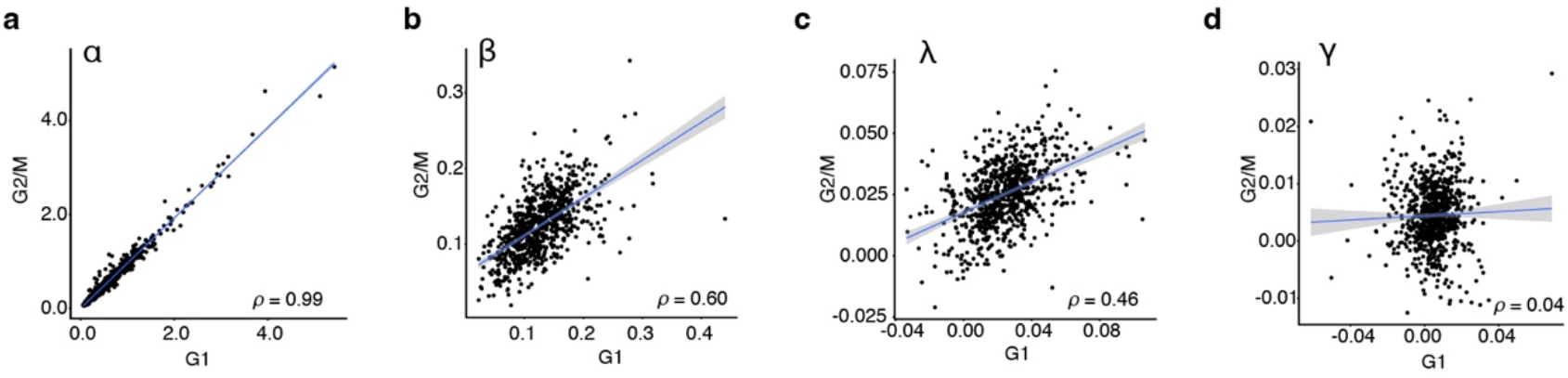
RNA kinetic parameter correlation in two cell-cycle stages. **a-d**, Examples of pairwise correlation in Fig. 3d, showing scatter plots of the relationships between G1 and G2/M. Pearson’s correlation coefficients from left to right: α (*ρ* = 0.99), β (*ρ* = 0.60), λ (*ρ* = 0.46), γ (*ρ* = 0.04).

**Extended data Fig. 6.**
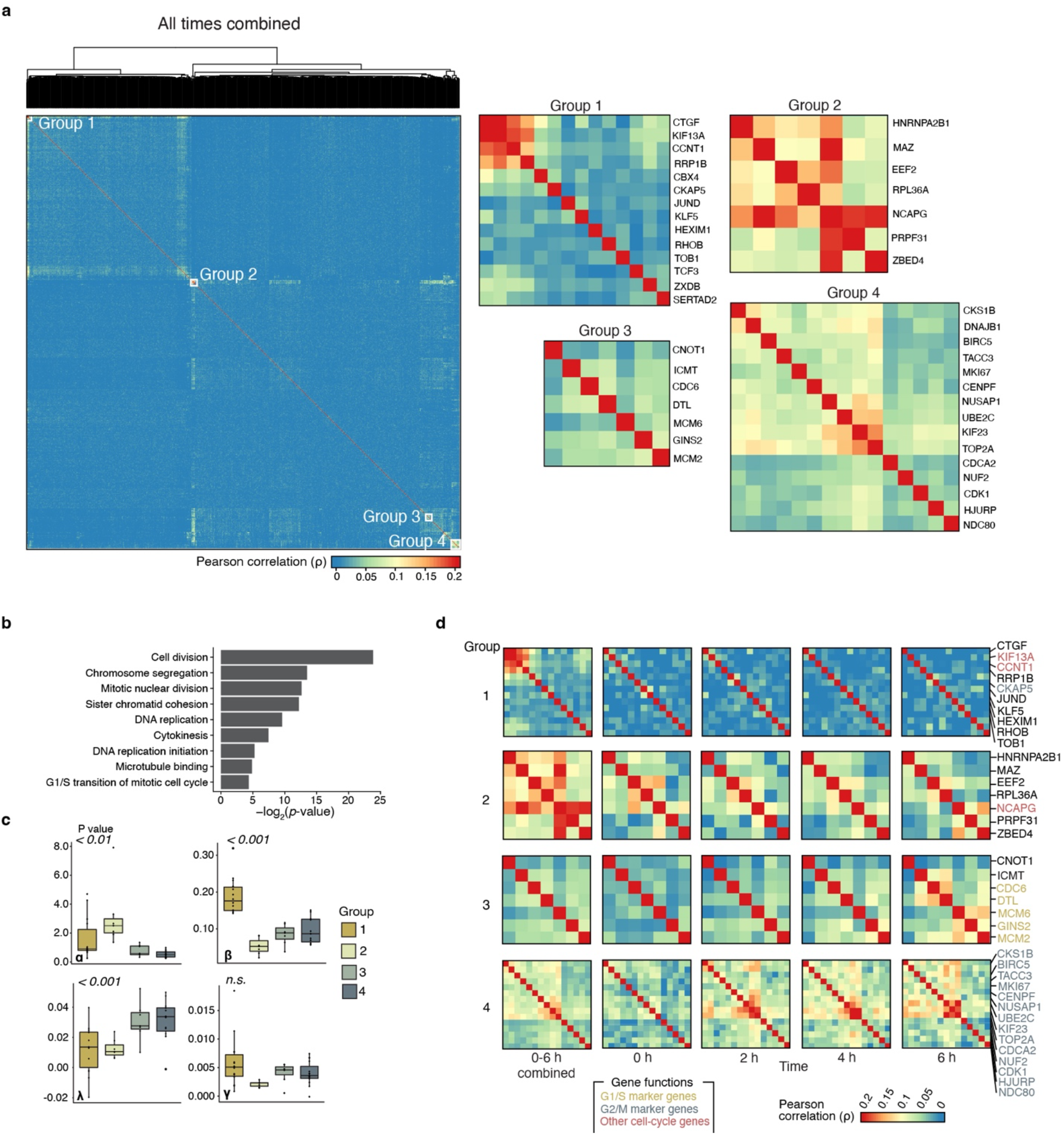
Single-cell RNA expression co-variation analysis. **a**, Heatmap depicting the pairwise correlation of 911 genes by TEMPOmap-measured single-cell RNA co-variation when combining four time points (0, 2, 4, 6 hrs chase), where the color indicates the value of Pearson correlation. Group 1-4 are highlighted for highly correlated gene modules (left) and zoomed-in (right). **b**, Pathway enrichment analysis result of genes in Group 1*-*4 from single-cell gene covariation heatmap in **a** using DAVID. **c**, Boxplots showing the distribution of six kinetic parameters of four gene groups (*n* = 14 genes in Group 1; *n* = 7 genes in Group 2, *n* = 7 genes in Group 3, *n* = 15 genes in Group 4). *P* values, one-way ANOVA test. Data shown as means, 25-75% quartiles and ranges. **d**, Heatmaps *were generated* showing matrices of the pairwise gene co-variation *in each of the* 0, 2, 4 and 6 hrs chase time points. Gene order along each matrix was the same and determined by the hierarchical clustering tree of the matrix combining the four time points in **a** (results were not shown). Zoom-in views of Group 1-4 from the co-variation heatmaps generated by gene expression in each time point, showing the correlation of RNA co-variation of each gene module across individual time points. Color in heatmaps indicates the value of Pearson correlation. G1/S and G2/M marker genes were annotated in each gene block.

**Extended data Fig. 7.**
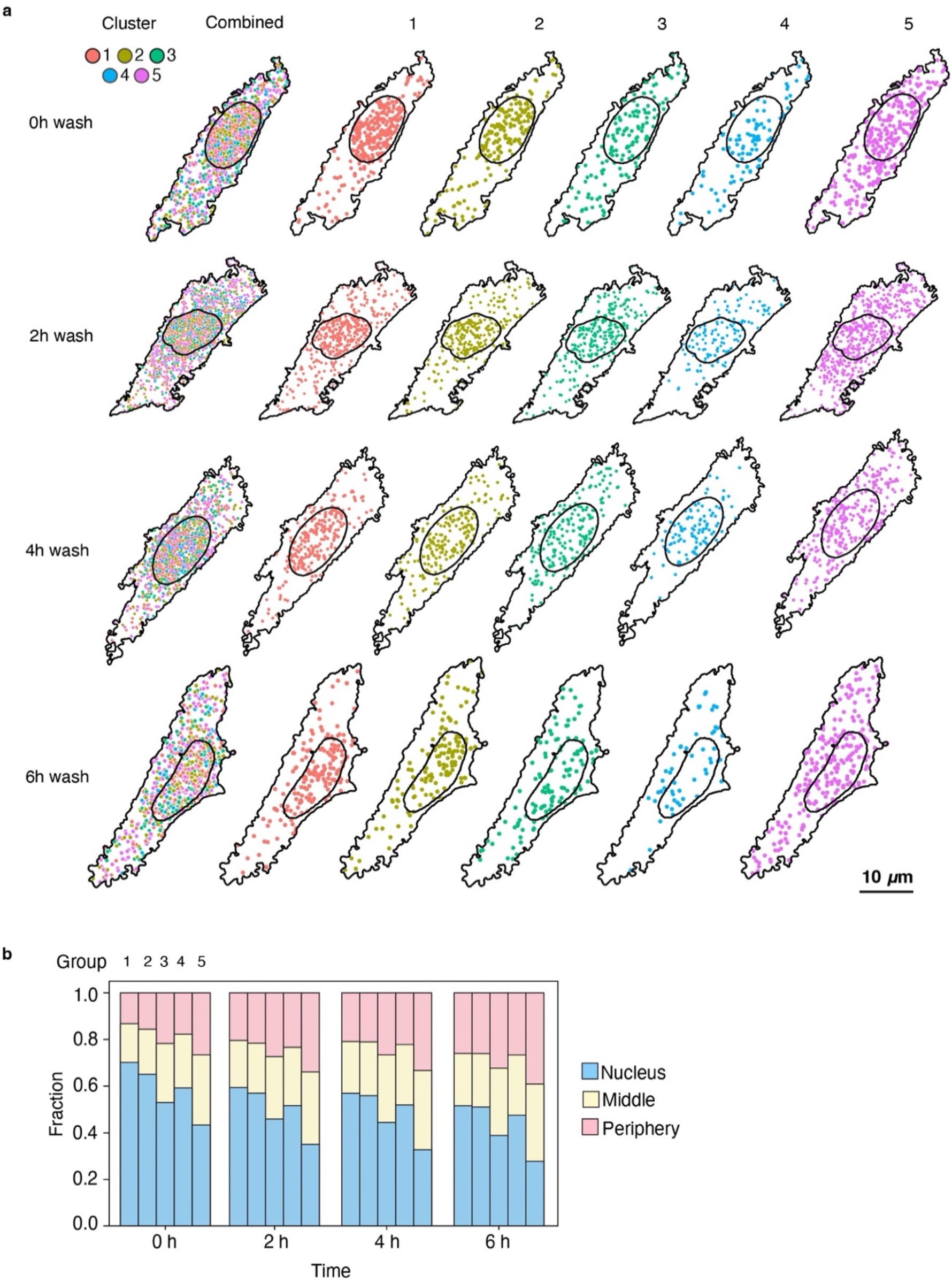
Visualization of gene clusters with different combination of RNA kinetic strategies. **a**, Visualization of all the amplicons in representative cells (combined, left) and separated by gene clusters (right) across pulse-chase time points. Scale bar: 10 *μ*m. Colors of amplicons indicate unique gene clusters. **b**, Boxplots showing the subcellular distribution of RNA reads over time in each kinetic cluster. Vertical lines indicate s.d. For each cluster, 0, 2, 4, 6 hrs chase have *n* = 1028, 910, 946, 542 cells, respectively.

**Extended data Fig. 8.**
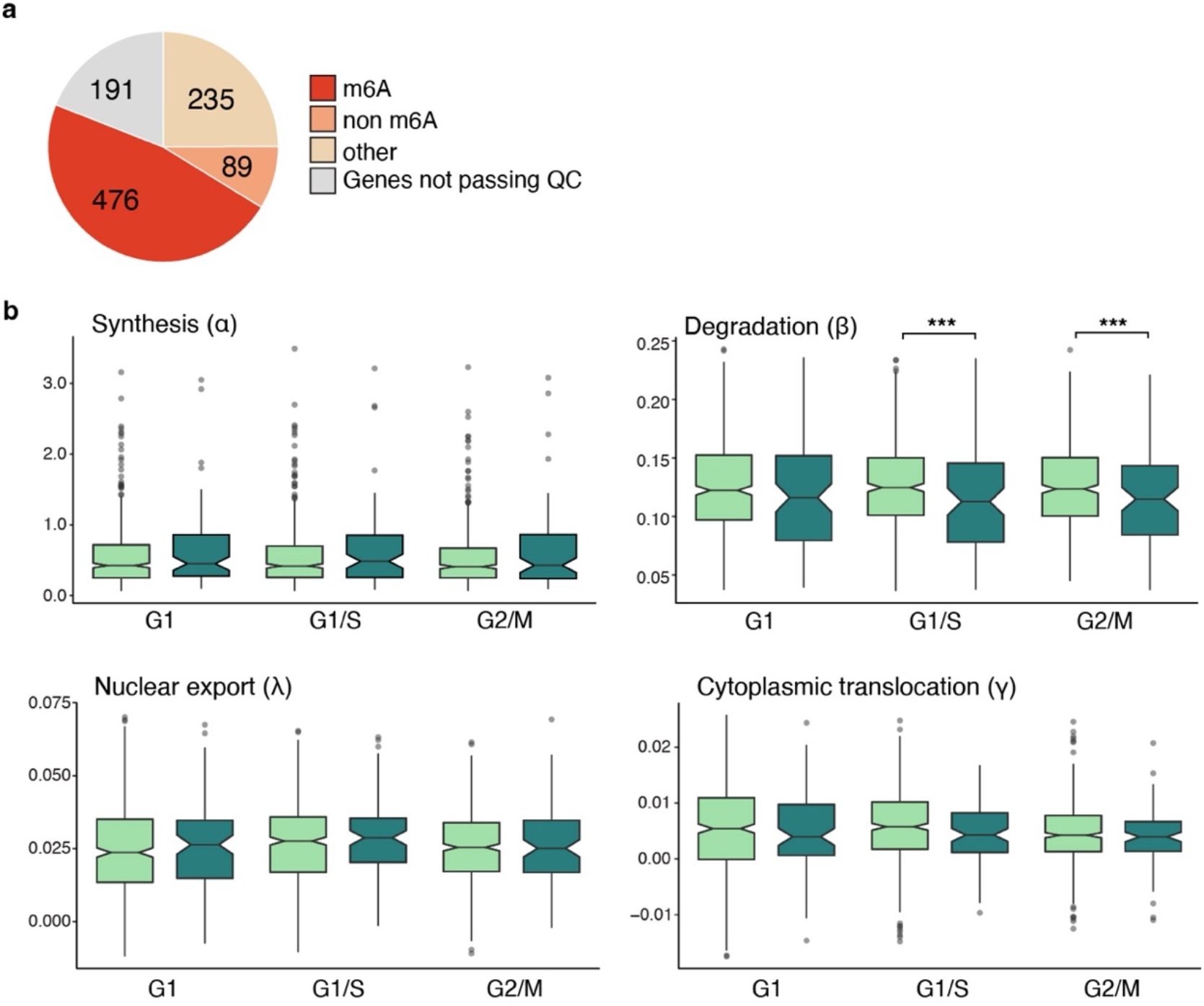
Subcellular RNA kinetics in the context of m^6^A post-transcriptional modification. **a**, Pie chart describing m^6^A-RNA methylation in the gene pool (see Methods). **c**, Boxplots comparing the four parameters estimated for m^6^A (*n* = 476 genes) and non-m^6^A RNAs (*n* = 89 genes) across three cell-cycle phases. *** *p*<0.01, Wilcoxin test. Data shown as means (notches), 25-75% quartiles (boxes) and ranges (vertical lines).

